# Blanding’s turtles (*Emydoidea blandingii*) as a reservoir for *Leptospira* spp

**DOI:** 10.1101/510149

**Authors:** Kelly E. Rockwell, Dan Thompson, Carol Maddox, Mark A. Mitchell

## Abstract

*Leptospira* spp. are re-emerging zoonotic pathogens. Previous research has found that Blanding’s turtles (*Emydoidea blandingii*) experimentally infected with *Leptospira interrogans* shed leptospires in their urine, suggesting that they could play a role in transmitting pathogen within an aquatic ecosystem. This study investigated whether a population of wild Blanding’s turtles known to be exposed to *Leptospira* spp. actively shed the pathogen under natural conditions. Blood samples were collected for serologic testing and to assess the health of the turtles. Free catch urine was collected for polymerase chain reaction (PCR) testing. All turtles were seropositive for *Leptospira* spp. and 73.5% (25/34) of the urine samples were PCR positive. All animals appeared clinically healthy and showed no apparent signs of disease. This study confirms that wild Blanding’s turtles can actively shed *Leptospira* spp. in their urine and suggests that they may play a role in the epidemiology of this disease in habitats in which they reside.

## Introduction

*Leptospira* spp. are important bacterial pathogens that can impact both humans and animals. As a result of urban sprawl, there is a concern that human exposure to this pathogen is increasing and that in certain regions of the world it is an important re-emerging zoonotic disease [1–3]. *Leptospira* spp. colonize the renal tubules of affected hosts and are subsequently shed in urine, leading to large accumulations of the pathogen in natural watersheds, lakes, and ponds. Animal and human exposure commonly occurs through contact with water contaminated with urine from infected animals or through infected animal sources via mucous membranes or damaged skin [1, 4]. Several domestic and wild mammals are known to serve as important reservoirs for serovars of *Leptospira* spp., including *L. kirschneri* serovar Grippotyphosa, L. *interrogans* serovar Canicola, and *L. borgpetersenii* serovar Hardjo [5, 6]. These three serovars, in addition to Pomona and Icterohaemorrhagiae, are also known to infect humans in the United States [7]. While research regarding the roles of wildlife in the epidemiology of *Leptospira* spp. grows, many species that could serve as important reservoirs remain undefined.

The role non-mammalian species play in the epidemiology of leptospirosis is not well understood. There has long been an interest in identifying the role chelonians serve in the dissemination of this disease; however, to date, much of this work has been experimental [6, 8–13]. Blanding’s turtles (*Emydoidea blandingii*) experimentally infected with *L. interrogans* serovar Pomona were found to become leptospiremic and shed the organism, suggesting that this species could play a role in the transmission of the pathogen in an aquatic ecosystem [10]. In addition, Andrews *et al.* [11] found a two-thirds higher rate of seroconversion in red-eared sliders (*Trachemys scripta elegans*) following infection compared with a terrestrial species of turtle (Eastern box turtle, *Terrapene carolina carolina*). However, this should not be unexpected as *Leptospira* sp. has a strong evolutionary association with water.

Blanding’s turtles (*Emydoidea blandingii*) are found throughout the upper Midwest and Northeast USA and are considered threatened or endangered throughout their range. Commonly found in marshes, prairie wetlands, wet sedge meadows, and shallow vegetated portions of lakes, their habitat is often threatened as a result of the pressures of urban sprawl. Because of their protected status, there are a number of groups performing long-term surveillance studies on these animals, which provides access to animals for sample collection. Protecting this species requires monitoring not only their general health, but also the environment in which they reside. Since this species has been found to experimentally shed *Leptospira* spp., it could play an important role in the epidemiology of the spirochete in aquatic habitats in which they reside. Because of the anthropogenic pressures on these systems, the dissemination of such a pathogen could have deleterious effects on the food web, ultimately leading to a negative impact on the protected Blanding’s turtle.

The purpose of this study was to determine if wild Blanding’s turtles actively shed *Leptospira* spp. under natural conditions. The hypotheses tested were that: 1) Blanding’s turtles would actively shed leptospires in their urine, and 2) that infected turtles would show no clinical signs of disease despite antibody recognition.

## Materials and methods

This cross-sectional study was performed in accordance to the regulations set forth by the University of Illinois Institutional Animal Care and Use Committee (IACUC protocol #14-087). Animals used for this study were part of an ongoing head-start program through the Forest Preserve District of DuPage County, IL, USA, and were trapped at five different locations within the preserve. Adult females that had been fitted with radio transmitters as part of the program were tracked for sampling. Any turtles opportunistically encountered during tracking were also sampled. Turtles were captured by hand and transferred to the Willowbrook Wildlife Center, Glen Ellyn, IL, USA where they were manually restrained for sampling.

Each turtle was identified by a passive integrated transponder tag and then received a physical exam. Morphometrics, including body weight, carapace length and width, plastron length and width, and body condition score, were collected from each animal. Sexing was based on externa sexual characteristics as well as presence of eggs in the coelomic cavity. Aging was based on size, with adults weighing >1000 grams and subadults weighing <1000 grams. Blood was collected from the jugular vein or subcarapacial sinus using a 3-ml syringe and a 22-gauge needle. A blood volume of less than one percent of each animal’s body weight was collected and placed in lithium heparin and anticoagulant-free microtainers (Becton-Dickinson, Franklin Lakes, NJ, USA) and processed for a complete blood count (CBC), biochemistry profile, and serologic testing. Blood smears were made immediately after the sample was collected. After the blood smears were made, the blood sample was centrifuged for 10 minutes at 1,500 × *g*. Serum was removed and stored in cryovials (Becton Dickinson, Franklin Lakes, NJ, USA). Free catch urine produced by inserting a swab into the animal’s cloaca was collected into a cryovial for real time polymerase chain reaction (rPCR) testing. Urine and serum samples were transported on frozen gel packs to the University of Illinois College of Veterinary Medicine’s Veterinary Diagnostic Laboratory for processing.

Complete blood cell counts and biochemistry profiles were used to assess health and physiologic status of the turtles. Air-dried blood smeared slides were stained with modified Wright-Giemsa stain (HemaTek Stain Pak, Bayer Corporation, Elkhart, IN, USA) and placed in dry storage boxes. White blood cell (WBC) estimates and differentials were performed manually by the same individual using standard techniques [4]. Briefly, an estimated white blood cell count was obtained by counting the number of white blood cells in 10 fields at 400x magnification, dividing that total number by 10, and multiplying the average by 2,000. Biochemistry testing was done using the VetScan and avian/reptile rotors (Abaxis Inc., Union City, CA, USA). The biochemistries measured in the turtles included: glucose, aspartate aminotransferase (AST), creatinine kinase (CK), total protein, albumin, globulin, calcium, phosphorus, sodium, potassium, and uric acid.

The *Leptospira* Microscopic Agglutination Test (MAT) was used to serologically screen the turtles. Serial two-fold dilutions of serum from 1:25 – 1:800 were evaluated against seven serovars commonly found in Illinois (*Leptospira interrogans* serovars: Autumnalis, Bratislava, Canicola, Icterohaemorrhagiae, Pomona; *Leptospira kirschneri* serovar Grippotyphosa; *Leptospira borgpetersenii* serovar Hardjo). The antigen was prepared from cultures grown in Probumin media (Millipore, Billeria, MA) and centrifuged at 349xg for ten minutes at room temperature to remove dead bacteria. Cultured bacteria are maintained based on proficiency requirements for American Association of Veterinary Laboratory Diagnostician (AAVLD) accredited laboratories. The supernatant was diluted 1:6 with sterile Phosphate Buffered Saline (PBS). Serum samples were then centrifuged at 349xg for one minute at room temperature to remove any red blood cells and lipids, and pipetted into a 96 well flat bottom plate. The sera were then diluted from 1:12.5 to 1:400 using 2-fold serial dilutions with PBS. End point sera dilutions were based on the National Veterinary Services Laboratories (NVSL) established guidelines for *Leptospira* spp. testing in wildlife. Fifty microliters of antigen were added to the fifty microliters of diluted sera. Plates were incubated at room temperature for 90-120 minutes and examined using dark field microscopy. The end point was determined as the last dilution exhibiting > 10% agglutination. A titer ≥ 1:25 was considered positive for wildlife (non-vaccinated titer) (Veterinary Diagnostics Laboratory Standard Operating Procedure, University of Illinois, Urbana, IL). The serovar that had the highest titer was considered causative. An animal was labeled as cross-reactive if it had reactions to multiple serovars at the same titer levels.

The cloacal swab with urine was brought to a 1.2 ml volume with 1X phosphate buffered saline and vortexed. The swab was removed and the suspension pelleted at 6,000 × g for 10 minutes. The pellet was resuspended in 110 ml of RLT buffer and the DNA was extracted utilizing the Qiagen® One-for-All Vet Kit BS96 and Vet 100 protocol on the BioSprint® Instrument. An rPCR was used to screen urine for the presence of *Leptospira* spp. DNA. The rPCR is based upon the primer sequences published by Smythe et al. [2] which generate an 87 bp amplicon confirmed by an internal probe 5’(FAM) CTCACCAAGGCGACGATCGGTAGC (TAMRA) 3’. The rPCR reaction mixture contained 1 OmniMix® bead, 41 μl nuclease free water, 1μl of each 10 μM primer, 2μl of 10 μM probe and 2.5 μl of extracted DNA sample. The rPCR assay was performed in a Cepheid Smart Cycler® with an initial cycle of 120 seconds at 95°C, then 50 cycles alternating 95°C for 15 seconds, and 60°C for 30 seconds. Results were recorded as positive (<38 Ct), weak positive (39-45 Ct with gel electrophoresis confirmation), or negative (0 Ct).

The 95% binomial confidence intervals (CI) were calculated for proportions. The distributions of the continuous CBC and biochemistry data were evaluated using the Shapiro-Wilk test, skewness, kurtosis, and box plots. Non-normal data were log transformed for parametric testing; however, they were reported by the median as it is the best estimate of central tendency for non-normal data. A single sample binomial test was used to determine if the sex ratio of this study population deviated from an expected proportion of 0.5. A Fisher exact or chi-square test was used to determine if there was any association between urine positivity and sex, age, or location. A follow-up logistic regression model was performed for any variables with a p<0.20 in the univariate analysis. Appropriate variables were entered into the model using a stepwise approach. The Hosmer-Lemeshow test was used to evaluate goodness-of-fit. A Mann-Whitney test was used to analyze differences between juvenile and adult titers. A Kruskal Wallis test was used to determine differences in titers for sex and the five different locations within the forest preserve. Independent samples t-tests were used to determine if there was a difference in CBC and biochemistry data by sex. Levene’s test was to assess equality of variance. Point biserial correlation coefficients were calculated to determine if there was an association between titer and the CBC and biochemistry data, as well as urine positivity and the bloodwork. SPSS 24.0 (IBM Statistics, Armonk, NY) was used to analyze the data, and a p<0.05 was used to determine statistical significance.

## Results

A total of 34 turtles were sampled: twenty-six females (76.5%), seven males (20.5%), and one juvenile (3%). The sex ratio of this population significantly (p=0.002) differed from a natural proportion of 0.5. This was not unexpected as the study subjects are part of an ongoing head start program. Each turtle was determined to be in good health and showed no apparent signs of infection or disease based on the physical examination, complete blood count, and biochemistry profile findings. [8]

All 34 turtles were seropositive for *Leptospira* spp. and 25 (73.5%, 95% confidence interval [CI]: 58.6-88.3) urine samples were PCR positive (Table 1). There was no significant difference in the likelihood of turtles shedding *Leptospira* spp. by sex (p=0.06), age (p=0.34), or location (p=0.06), although sex and location approached significance. None of the variables were included in a final logistic regression model. The most common serotypes found in this population of turtles were Hardjo (n=13; 38.2%, 95% CI: 21.9-54.5), Autumnalis (n=9; 26.5%, 95% CI: 11.7-41.3), and Icterohaemorrhagiae (n=1; 2.9%, 95% CI: 0.1-8.6), with 11 (32.4%, 95% CI: 16.7-48.1) animals having mixed serotypes (Table 1). There was no significant difference in serotypes by age (*χ*^2^=3.4, p=0.33), sex (*χ*^2^=4.6, p=0.20), or location (*χ*^2^=6.2, p=0.72).

**Table 1.** Serology and RT-PCR results for Blanding’s turtles from DuPage County Forest Preserve.

| Turtle no. | Sex | Age | Location | PCR +/- | Immunoreactive serovar(s) | Highest titer |
| --- | --- | --- | --- | --- | --- | --- |
| 1 | F | Adult | 1 | - | H | 1:200 |
| 2 | F | Adult | 5 | - | A, I | 1:100 |
| 3 | F | Adult | 2 | + | A, B, P | 1:200 |
| 4 | F | Adult | 5 | + | A | 1:200 |
| 5 | F | Adult | 2 | - | A | 1:200 |
| 6 | F | Adult | 2 | + | A | 1:200 |
| 7 | F | Adult | 2 | + | A | 1:400 |
| 8 | F | Adult | 2 | + | H | 1:200 |
| 9 | M | Adult | 5 | + | A | 1:100 |
| 10 | M | Adult | 5 | + | H | 1:100 |
| 11 | F | Adult | 3 | - | H | 1:800 |
| 12 | F | Adult | 2 | + | H | 1:100 |
| 13 | M | Adult | 4 | + | I | 1:100 |
| 14 | M | Adult | 4 | + | A, C, H, I, P | 1:50 |
| 15 | M | Adult | 4 | + | A, C, H, I | 1:50 |
| 16 | F | Subadult | 5 | + | A | 1:400 |
| 17 | F | Adult | 1 | - | A, I | 1:200 |
| 18 | F | Adult | 2 | + | H, I | 1:50 |
| 19 | F | Adult | 2 | + | A, I | 1:100 |
| 20 | F | Adult | 2 | - | A, H | 1:100 |
| 21 | F | Adult | 2 | - | A | 1:800 |
| 22 | F | Adult | 2 | - | A, H | 1:50 |
| 23 | F | Adult | 2 | + | H | 1:100 |
| 24 | F | Adult | 2 | + | H | 1:400 |
| 25 | F | Adult | 2 | - | H | 1:200 |
| 26 | F | Adult | 2 | + | H | 1:400 |
| 27 | F | Adult | 4 | + | H | 1:100 |
| 28 | F | Adult | 1 | + | H | 1:100 |
| 29 | F | Adult | 4 | + | H, I | 1:200 |
| 30 | M | Adult | 4 | + | H | 1:400 |
| 31 | F | Adult | 4 | + | A | 1:100 |
| 32 | F | Adult | 4 | + | H | 1:100 |
| 33 | M | Subadult | 2 | + | A, H, I | 1:50 |
| 34 | UNK | Juvenile | 2 | + | A | 1:200 |

Sex: F=female, M=male, UNK=unknown; Immunoreactive serovars are as listed: A = Autumnalis, B = Bratislava, C = Canicola, I = Icterohaemorrhagiae, H = Hardjo, P = Pomona

There were significant differences in calcium (t=−9.438, p=0.0001), albumin (t=−2.088, p=0.046), and glucose (t=3.264, p=0.003) by sex (Table 2); none of the other biochemistry data differed by sex (phosphorus, t=−1.128, p=0.268; total protein, t=−1.194, p=0.241; globulins, t=−0.540, p=0.593; potassium, t=−0.638, p=0.528; sodium, t=1.367, p=0.181; creatinine kinase, t=−0.193, p=0.848; aspartate aminotransferase, t=−1.041, p=0.306; uric acid, t=−0.046, p=0.964). There were no differences in any of the CBC findings by sex (WBC, t=1.181, p=0.246; heterophils, t=−0.196, p=0.846; lymphocytes, t=0.259, p=0.797; monocytes, t=−0.204, p=0.839; eosinophils, t=− 0.550, p=0.586; basophils, t=−0.399, p=0.693).

**Table 2.** Biochemistry differences by sex.

| *Parameter* | *Sex* | *Median* | *25-75%* | *Minimum-Maximum* |
| --- | --- | --- | --- | --- |
| Albumin | Male | 1.45 | 1.25-1.6 | 1.10-1.60 |
| (g/dL) | Female | 1.70 | 1.50-1.90 | 1.10-2.50 |
| Glucose | Male | 80.5 | 70.5-117.75 | 60.0-120.0 |
| (mg/dL) | Female | 55.0 | 50.0-71.0 | 46.0-110.0 |
| Calcium | Male | 11.5 | 10.77-11.97 | 10.4-12.8 |
| (mg/dL) | Female | 20.0 | 15.95-20.0 | 10.9-20.0 |

There were no significant correlations found between titer and the CBCs or the biochemistries (WBC, r=−0.008, p=0.962; heterophils, r=0.218, p=0.216; lymphocytes, r=−0.216, p=0.219; monocytes, r=0.079, p=0.655; eosinophils, r=0.180, p=0.308; basophils, r=−0.147, p=0.406; total protein, r=0.027, p=0.880; albumin, r=0.031, p=0.866; globulin, r=−0.08, p=0.663, potassium, r=0.053, p=0.766; glucose, r=0.026, p−0.882; sodium, r=−0.306, p=0.08; phosphorus, r=−0.045, p=0.802; calcium, r=0.228, p=0.194; creatinine kinase, r=0.311, p=0.078; aspartate aminotransferase, r=0.145, p=0.415; uric acid, r=−0.153, p=0.485). There were also no significant correlations found between urine PCR status and the CBCs or the biochemistries (WBC, r=0.079, p=0.657; heterophils, r=0.208, p=0.239; lymphocytes, r=−0.150, p=0.396; monocytes, r=−0.154, p=0.383; eosinophils, r=−0.186, p=0.292; basophils, r=−0.093, p=0.601; total protein, r=−0.159, p=0.368; albumin, r=−0.108, p=0.556; globulin, r=−0.280, p=0.120, potassium, r=−0.247, p=0.160; glucose, r=0.310, p=0.074; sodium, r=0.103, p=0.561; phosphorus, r=0.011, p=0.950; calcium, r=−0.323, p=0.062; creatinine kinase, r=−0.040. p=0.826; aspartate aminotransferase, r=−0.109, p=0.541; uric acid, r=0.087, p=0.694).

## Discussion

Leptospirosis is a reemerging disease with a cosmopolitan distribution [1–3]. To reduce or eliminate the risk of exposure to this pathogen, it is essential that relevant reservoir hosts be identified [6]. Historically, positive MAT titers have been used to determine whether wild reptiles serve as reservoirs for this important disease [14–19]; however, the serologic assay is limited in its scope, as it notes exposure but not active infection without serial samples and a rise in titer. Because of this, there is a strong argument for PCR testing of reptiles to truly characterize them as reservoirs [17]. In South America, Oliveira *et al* reported a 16.7% (11/66) prevalence of *Leptospira* spp. from PCR samples collected from the stomach and cloaca of wild *Phrynops geoffroanus* in an urban environment in Brazil [14], and Biscola *et al* identified the serotype Interrogans from PCR samples of the liver and kidney from a wild *Bothrops pauloensis* [15]. In Africa, a wild *Crotaphopeltis hotamboeia* was positive for *Leptospira* spp. from PCR samples of kidney tissue [18]. The findings from the current study represent the first time *Leptospira* spp. has been confirmed using rPCR from ante-mortem samples collected from a wild aquatic chelonian species in the United States. The results of this study suggest that Blanding’s turtles may be an important reservoir in the aquatic ecosystems that they reside. The high prevalence of shedding, in combination with an absence of obvious disease, suggests that these animals may carry and disseminate the pathogen. It is also possible that they may serve as an over wintering source of the pathogen.

It is interesting to note that multiple serovars were identified in these animals, suggesting that they can serve as reservoirs to organisms found to infect a variety of higher vertebrates, including humans. In a previous study evaluating *Leptospira* serovars in this population of turtles, the authors found a similar high seroprevalence (93.5%, 29/31 animals); the two seronegative animals in that study seroconverted in this study [8]. In the first study, Icterohaemorrhagiae was the most common serovar (58%, 17/29); however, in this study, Icterohaemorrhagiae was only found in a single turtle (2.9%, 1/34) [8]. In the current study, Hardjo (38.2%, 13/34) and Autumnalis (26.5%, 9/34) were the most common serovars; these specific serovars were not identified in the first study [8]. One explanation for this difference may be that mixed serovars were noted in 20.7% (6/30) and 32.4% (11/34) of the turtles, respectively, and that cross-reactions between serovars are common. Future studies evaluating the *Leptospira* DNA may be necessary to better characterize the source and movement of this pathogen in these turtles and their aquatic ecosystem.

A 1999 study assessing the serological titers of infectious organisms in raccoons (*Procyon lotor*) in west-central Illinois (USA) found that of 459 raccoons sampled, 222 (48%) were seropositive for *L. interrogans* [6]. In the current study, 70.6% of the Blanding’s turtles were PCR positive for *Leptospira* spp.; a high prevalence that is comparable to a wild mammalian reservoir within a similar environment. PCR samples were not sequenced in the current study, but considering the variety of serovars found in these turtles, identifying the type(s) present in each animal would be important future work to fully understand the role these turtles play in the epidemiology of this reemerging disease.

It is important to recognize that Blanding’s turtles are a protected species. Habitat fragmentation and loss have led to diminished aquatic habitat for these animals, leading to higher densities of the turtles. This can lead to increased competition and reduced survival. The results of this study suggest it may also lead to an increased exposure to pathogens, as the presence of leptospiral DNA was detected in the urine of nearly three-quarters of the turtles. It is possible that with future environmental stressors and higher pathogen loads, these turtles could become susceptible to *Leptospira* spp. infections and begin to exhibit clinical signs. As the environmental load of *Leptospira* spp. increases and the interface between wildlife habitats and developed land diminishes, domestic animals are at greater risk of exposure and infection, which can lead to more profound effects [4, 6]. Fortunately, at this time, the turtles appear to be unaffected based on their normal examination findings and blood work. The only significant differences in blood work were associated with calcium, albumin and glucose between the sexes. The higher calcium and albumin concentrations were not unexpected because the turtles were sampled during the breeding season and these biochemistries increase in females during oviposition. The lower glucose noted in the females was also presumed to be associated with the breeding season, as females become inappetent during oviposition [8].

The results of this study strongly suggest that more work is needed to further characterize the role of Blanding’s turtles, as well as other aquatic turtles, in the epidemiology of *Leptospira* spp. At the same time, it is important to consider the influences of such a disease on the turtles too, especially as the pathogen density increases in these restricted aquatic habitats. Veterinarians and biologists will need to work together to protect this important species and ensure it has a successful future.

